# Sorting and cultivation of *Faecalibacterium prausnitzii* from fecal samples using flow cytometry in anaerobic conditions

**DOI:** 10.1101/2020.03.25.007047

**Authors:** Samuel Bellais, Mélanie Nehlich, Aurore Duquenoy, Maryne Ania, Ger van den Engh, Jan Baijer, Ilia Belotserkovsky, Vincent Thomas

## Abstract

**Background:** There is a growing interest in using gut commensal bacteria as ‘next generation’ probiotics. However, this approach is still hampered by the fact that there are few or no strains available for specific species that are difficult to cultivate. Our objective was therefore to adapt flow cytometry and cell sorting to be able to detect, separate, isolate and cultivate new strains of Extremely Oxygen Sensitive (EOS) species from fecal material, focusing on *Faecalibacterium prausnitzii* as a proof-of-concept.

**Results:** A BD Influx^®^ cell sorter was equipped with a glovebox that covers the sorting area. This box is flushed with nitrogen to deplete oxygen in the enclosure. Several non-specific staining methods including Wheat Germ Agglutinin (WGA), Vancomycin BODIPY™ and LIVE/DEAD BacLight were evaluated with three different strains of the EOS species *F. prausnitzii*. In parallel, we generated polyclonal antibodies directed against this species by immunizing rabbits with heat-inactivated bacteria. Anaerobic conditions were maintained during the full process, resulting in only minor viability loss during sorting and culture of unstained *F. prausnitzii* reference strains. In addition, staining solutions did not severely impact bacterial viability while allowing discrimination between groups of strains. Efficient detection was achieved using polyclonal antibodies directed against heat-fixed bacteria. Finally, we were able to detect, isolate and cultivate a variety of *F. prausnitzii* strains from healthy volunteer’s fecal samples using WGA staining and antibodies. These strains present markedly different phenotypes, thus confirming the heterogeneity of the species.

**Conclusions:** Cell-sorting in anaerobic conditions is a promising tool for the study of fecal microbiota. It gives the opportunity to quickly analyze microbial populations and to sort strains of interest using specific antibodies, thus opening new avenues for targeted culturomics experiments.

## Background

The compositional analysis of commensal bacterial populations collected from a variety of clinical samples has been recently made possible with the availability of Next Generation Sequencing (NGS) technologies. In particular, fecal microbiota analysis is now widely used as a proxy of gut microbiota analysis. In this case, 16S repertoire or whole genome shotgun analysis conducted with cohorts of patients vs controls highlighted the positive or negative association of a variety of commensal bacterial species or group of species with a number of pathological conditions. In the case of negative association, *i.e*. commensals showing decreased abundancy in pathological conditions, there is a growing interest in using cultivated, well-characterized strains to complement deficiencies in the gut microbiota. The term ‘next-generation probiotics’ (NGP) is now widely used to describe these commensal species of potential health interest. The Extremely Oxygen Sensitive (EOS) species *Faecalibacterium prausnitzii* and the mucin foraging species *Akkermansia muciniphila* are the most extensively described examples of such NGP, with live bacteria or bacteria-derived products being developed by private companies for specific indications such as obesity or Inflammatory Bowel Disease (IBD).

Knowing that specific biological properties, including host beneficial properties, can vary significantly from one bacterial strain to another, there is a major interest in building collections of commensal strains corresponding to one species of interest that has been identified based on NGS. However, this approach is still hampered by the fact that there are usually few or even no strains available for a number of commensal species. This can be due to the fact that the targeted species have specific nutritional requirements rendering them difficult to cultivate in synthetic media. Extreme sensitivity to oxygen exposure or under-representation of the target species in the whole bacterial population can also be a factor. Due to these reasons, retrieving target species from clinical samples (usually fecal material) can be difficult.

In this context, we believe that flow cytometry (FCM) coupled with cell-sorting has the potential to circumvent most if not all these limitations. With constantly increasing technological performances, it can be used for bacterial or even viral cell populations’ analysis with or without subsequent sorting. With the objective to use FCM and cell-sorting to analyze, sort and cultivate bacterial species of interest from fecal samples, we adapted a cell sorter and associated workflow to conduct sorting experiments in complete anaerobic conditions. We then evaluated the impact of sorting as well as non-specific and specific staining methods on the viability of several representative strains of the EOS species *F. prausnitzii*. Specific targeting was finally used to sort and cultivate new *F. prausnitzii* strains from fecal samples, demonstrating the potential of the approach.

## Results

### Validation of Gram staining and sorting

First, we evaluated the sensitivity and efficiency of bacteria sorting using a Vancomycin molecule, which binds to the cell wall of the Gram positive bacteria, coupled to a BODIPY fluorophore enabling its detection by FCM. For this end, *E. coli* and *E. faecalis* bacteria (separate or in a 1:1 mix) were labeled with Vancomycin BODIPY and sorted onto the URI*Select*™ 4 plates Colonies sorted based on Vancomycin BODIPY™-FL staining were observed after overnight incubation at 37°C (Fig. 1). All colonies obtained from the Vancomycin BODIPY™-FL-negative gate were pink, corresponding to *E. coli*, whereas all colonies obtained from the Vancomycin BODIPY™-FL-positive gate were turquoise-blue, corresponding to E. faecalis. Colonies grown from spots seeded with one Vancomycin BODIPY™-FL-negative event plus one Vancomycin BODIPY™-FL-positive event appeared purple and were easily distinguishable from single-species colonies. In order to determine if only one bacterium was deposited on each spot, bacteria randomly selected from a 1:1 mixture of *E. coli*/*E. faecalis* were sorted onto URI*Select*™ 4 agar plates. 174 colonies were observed out of 180 seeded events (recovery rate: 95%), with no purple spots being detected, indicating that a single bacterium was effectively deposited at each position.

**Fig. 1.**
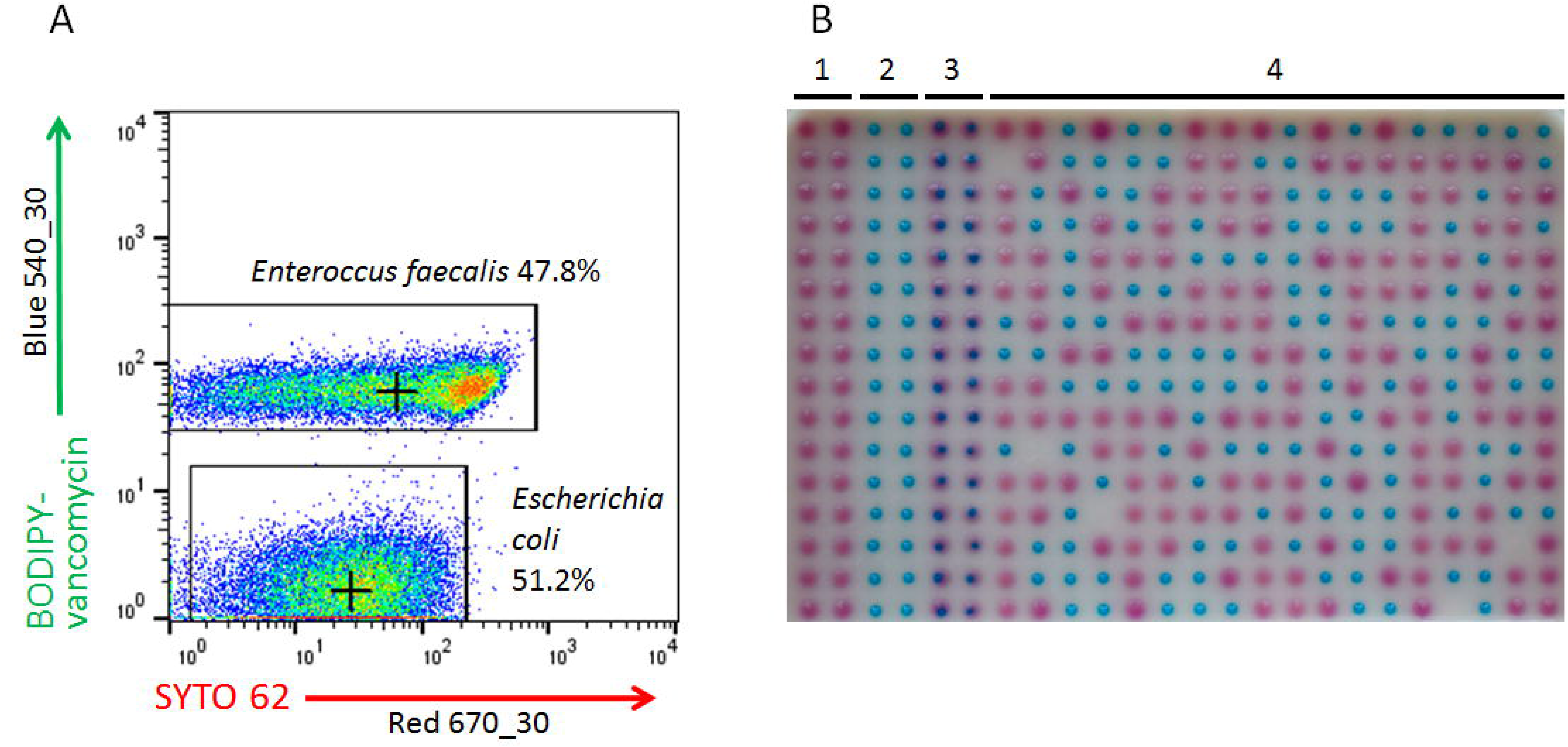
Validation of Gram staining and sorting precision. A 50/50 mix of Gram positive (*E. faecalis*) and Gram-negative (*E. coli*) bacteria was double-stained with Vancomycin BODIPY™-FL, a probe that binds to peptidoglycan, and cell-permeant nucleic acid stain SYTO 62 (panel A). Bacteria were then sorted on a URI4™ select agar plate according to Vancomycin BODIPY™-FL stain: unstained bacteria for B1, stained bacteria for B2, and one of each (stained/unstained) for B3. Bacteria were then randomly sorted for region B4. *E. coli* appear as pink colonies, *E. faecalis* as blue colonies.

## Staining of *F. prausnitzii* reference strains

Antibodies generated by vaccinating rabbits with heat-inactivated bacteria were very efficient at detecting both ATCC strains used for vaccination, with approximately 90% of the bacteria displaying strong fluorescence signal after labelling (Fig. 2b). Conversely, the strain A2-165 was not detected at all by the antibodies (<1% of the gated events). The reverse situation was observed for WGA staining, with strain A2-165 being efficiently stained by the lectin (>90% of the gated events) whereas strains ATCC 27766 and ATCC 27768 were less efficiently stained (10.9 and 3.8% of the gated events, respectively) (Fig. 2c). A variable intensity autofluorescence was observed at 540 nm for the three strains upon excitation with the 488 nm laser (Fig. 2d). Once stained with the Vancomycin BODIPY™-FL dye, bacteria from the 3 reference strains did not display significantly higher levels of fluorescence compared to unstained bacteria, thus rendering the use of this staining useless for the selection of *F. prausnitzii* from complex populations (data not shown).

**Fig. 2.**
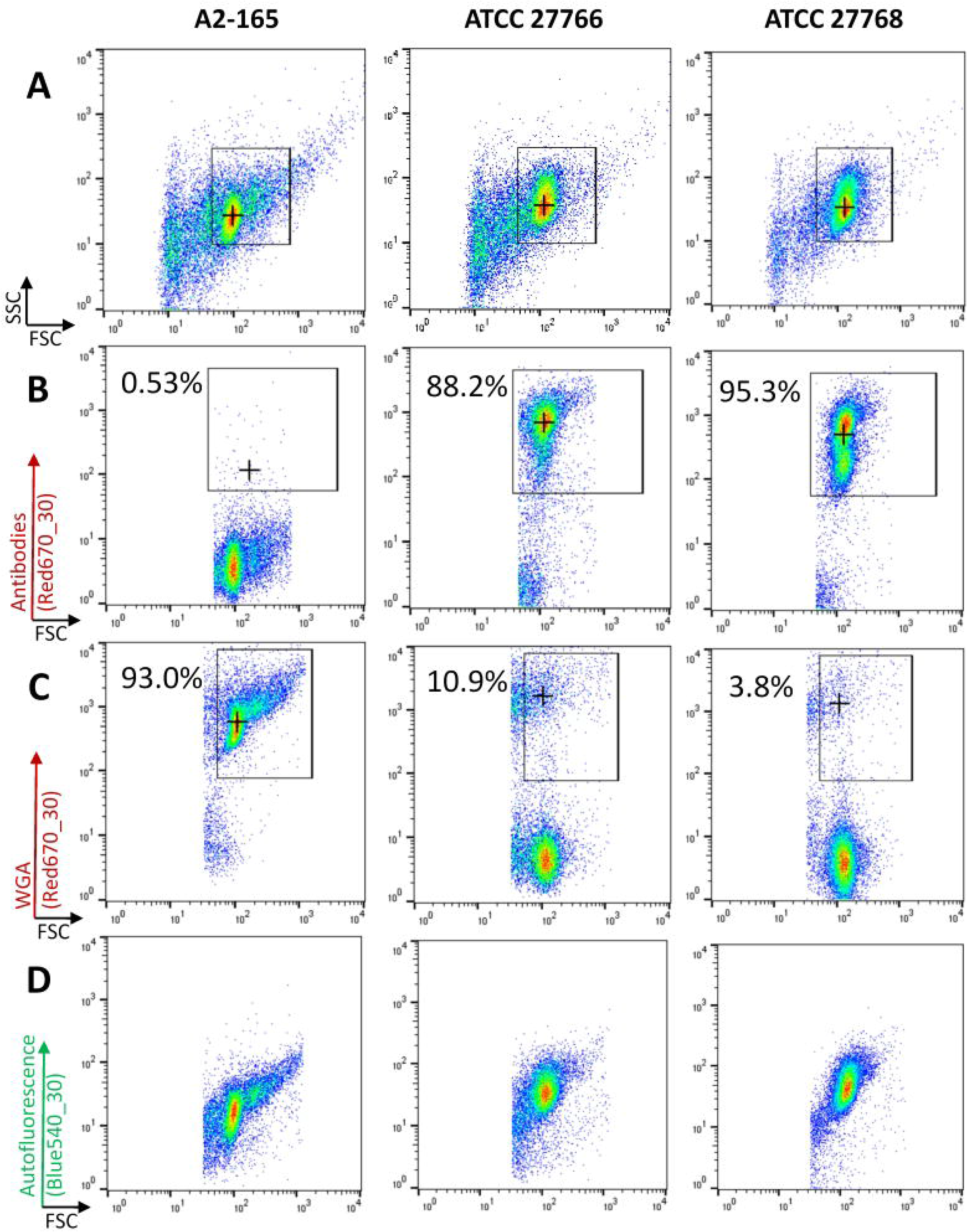
Staining of *F. prausnitzii* reference strains. Fresh cultures of the 3 F.prausnitzii reference strains: DSM-17677 (A2-165), ATCC-27766 and ATCC-27768 were analyzed after staining with: polyclonal antibodies generated with both ATCC strains (B), or Wheat Germ Agglutinin (C). Natural auto-fluorescence of unstained bacteria was also observed at 540nm (D). These measurements were made only with gated events corresponding to the vast majority of bacteria in the suspensions (A).

## Impact of staining and anaerobic sorting on *F. prausnitzii* cultivability

We compared two conditions to evaluate the impact of atmospheric oxygen on bacterial cultivability: (i) staining and sorting performed in anaerobic conditions, (ii) staining in anaerobic conditions followed by sorting in atmospheric conditions. Durations of the staining (including washing steps) and of the sorting were normalized at 30 min for each step. Paraffin oil was used to cover samples in both conditions. Overall recovery of unlabeled *F. prausnitzii* after sorting in anaerobic conditions was approximately 20% for the three tested strains, while no colony could be observed if the sorting step was performed in atmospheric conditions (Fig. 3). We did not observe significant impact of any of the tested staining (SYTO 9 in the LIVE/DEAD kit, WGA-647 and Vancomycin BODIPY) on recovery efficiency compared to unlabeled bacteria (Fig. 4). Several colonies were still observed after anaerobic sorting and cultivation of the propidium iodide (PI)-stained fraction, corresponding to approx. 0.1 to 1% of cultivable bacteria in this fraction (Fig. 4).

**Fig. 3.**
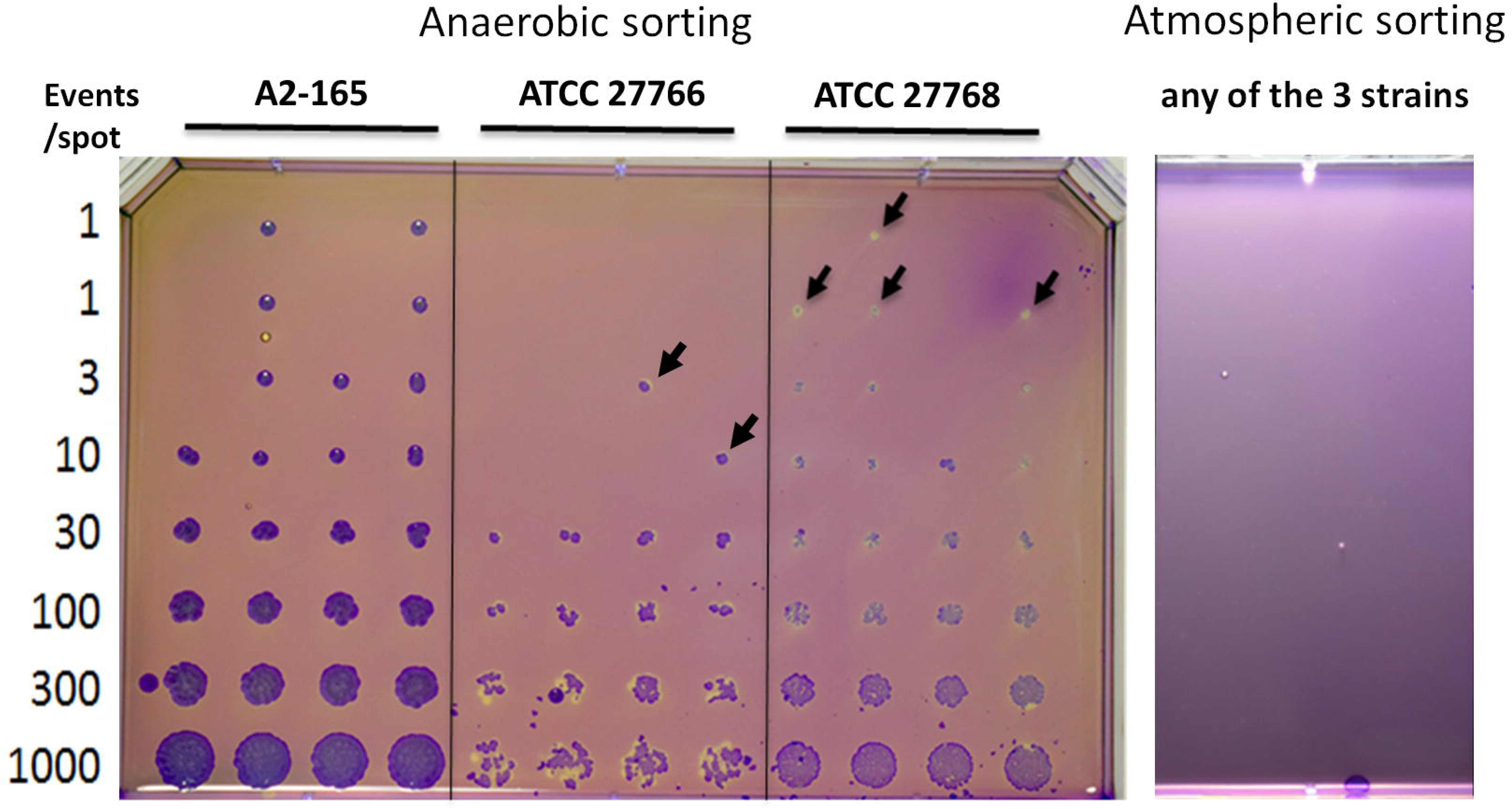
Impact of sorting in anaerobic or atmospheric conditions on *F. prausnitzii*’s recovery. Fresh cultures of the 3 F.prausnitzii reference strains were sorted on mGAM plates in anaerobic (left) or atmospheric (right) conditions. Increasing (1 to 1,000) amounts of bacteria were sorted on each spot. Colonies were stained with crystal violet for better visualization of tiny colonies (black arrows). Experiments were repeated at least 3 times.

**Fig. 4.**
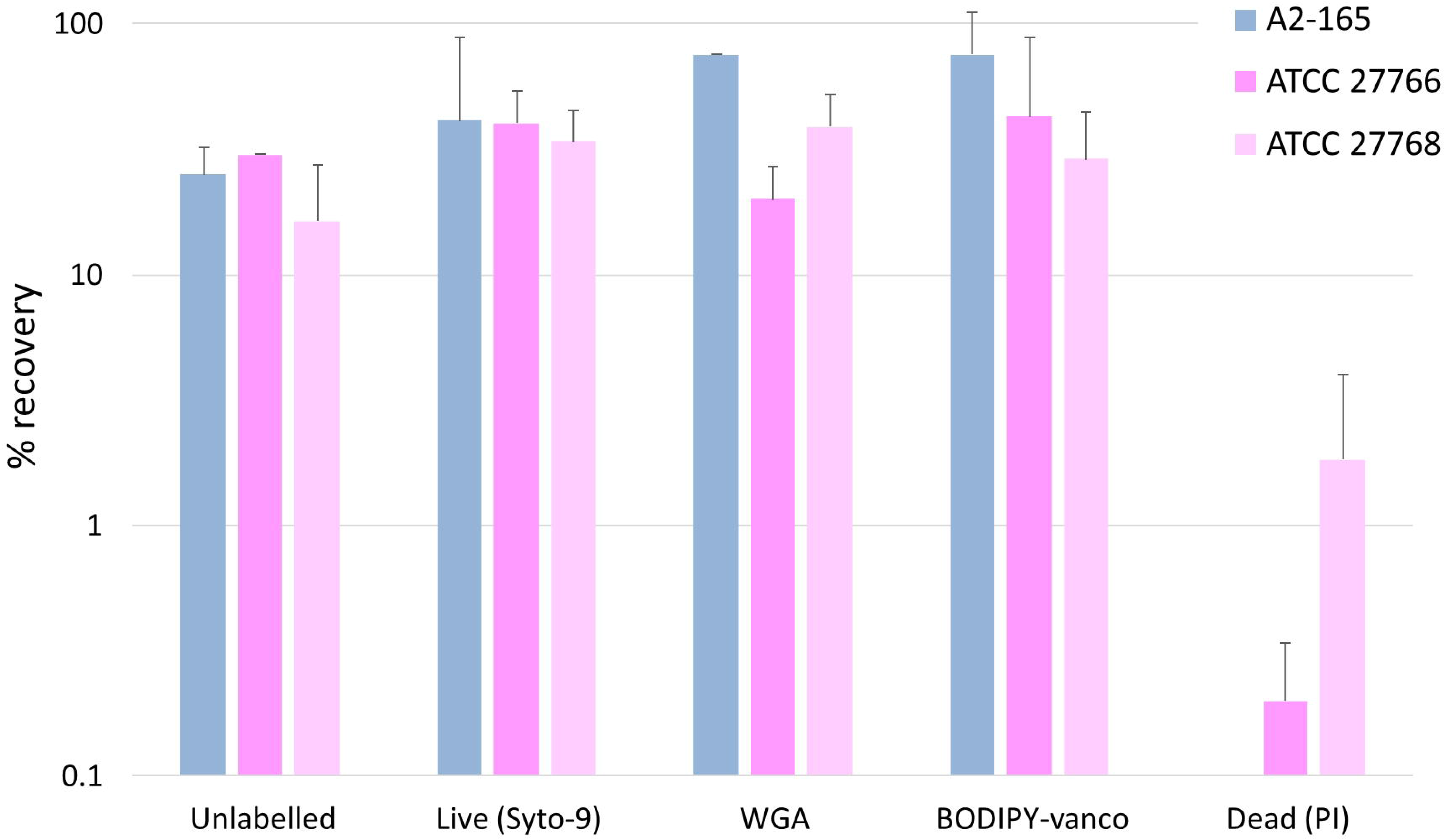
Proportions of bacteria recovered in culture after various staining. Fresh cultures of the 3 *F.prausnitzii* reference strains were stained with syto 9, WGA, Vancomycin BODIPY™-FL or propidium iodide, and then stained bacteria were sorted in increasing amounts on mGAM plates to calculate the percentage of recovery.

## Sorting and cultivation of new *F. prausnitzii* strains from fecal material

Based on the results observed with pure cultures of *F. prausnitzii* collection strains, we chose to associate LIVE/DEAD staining with a double staining with specific polyclonal antibodies and WGA-CF405M fluorescent lectin to perform sorting experiments from fecal material. In a first series of experiments with fecal samples collected from 5 healthy volunteers, live (*i.e*. SYTO 9 stained and IP negative) bacteria ranged from 13.3 to 46.1%, among which proportions of antibody-labelled bacteria ranged from 0.1 to 18.6% whereas proportions of WGA-stained events ranged from 0.6 to 2.3% (Table 1 & Fig. 5). Bacterial colonies cultivated from antibody and WGA-labeled fractions were then screened using *F. prausnitzii*-specific primers, resulting in proportions of PCR-positive colonies ranging from 0 to 53.8% in the antibody-stained fractions and 1.2 to 23.1% in the WGA-stained fractions (Table 1). In a second series of experiments conducted with 5 additional fecal samples anonymously collected from 5 healthy volunteers, the proportions of live bacteria ranged from 39.1 to 65.9 % and those of bacteria labeled with the polyclonal antibody varied from 0.1 to 4.1%, whereas the proportions of WGA-stained bacteria ranged from 1.9 to 8.3% (Table 2). In this second series, the events that were double stained by WGA and the antibody, ranging from 0.3 to 1.9% of the live bacteria, were also sorted for cultivation. For every fecal sample, 120 events corresponding to each of the 3 regions were sorted and cultivated on mGAM plates, resulting in variable numbers of colonies depending on the fecal samples and on the gated regions (Table 2). Colonies presenting morphologies compatible with *F. prausnitzii* (*i.e*. exclusion of large colonies that appeared in less than 48 h) were screened with species-specific primers, resulting in 9 PCR-positive colonies isolated from 3 different donors in the antibody-gated events, 8 PCR-positive colonies isolated from 5 different donors in the WGA-gated events and 1 PCR-positive colony isolated from 1 donor in the WGA + antibody-gated events (Table 2). These colonies were sub-cultivated for DNA extraction and then stored at −80°C in mGAM + 30% glycerol for long term preservation.

**Fig. 5.**
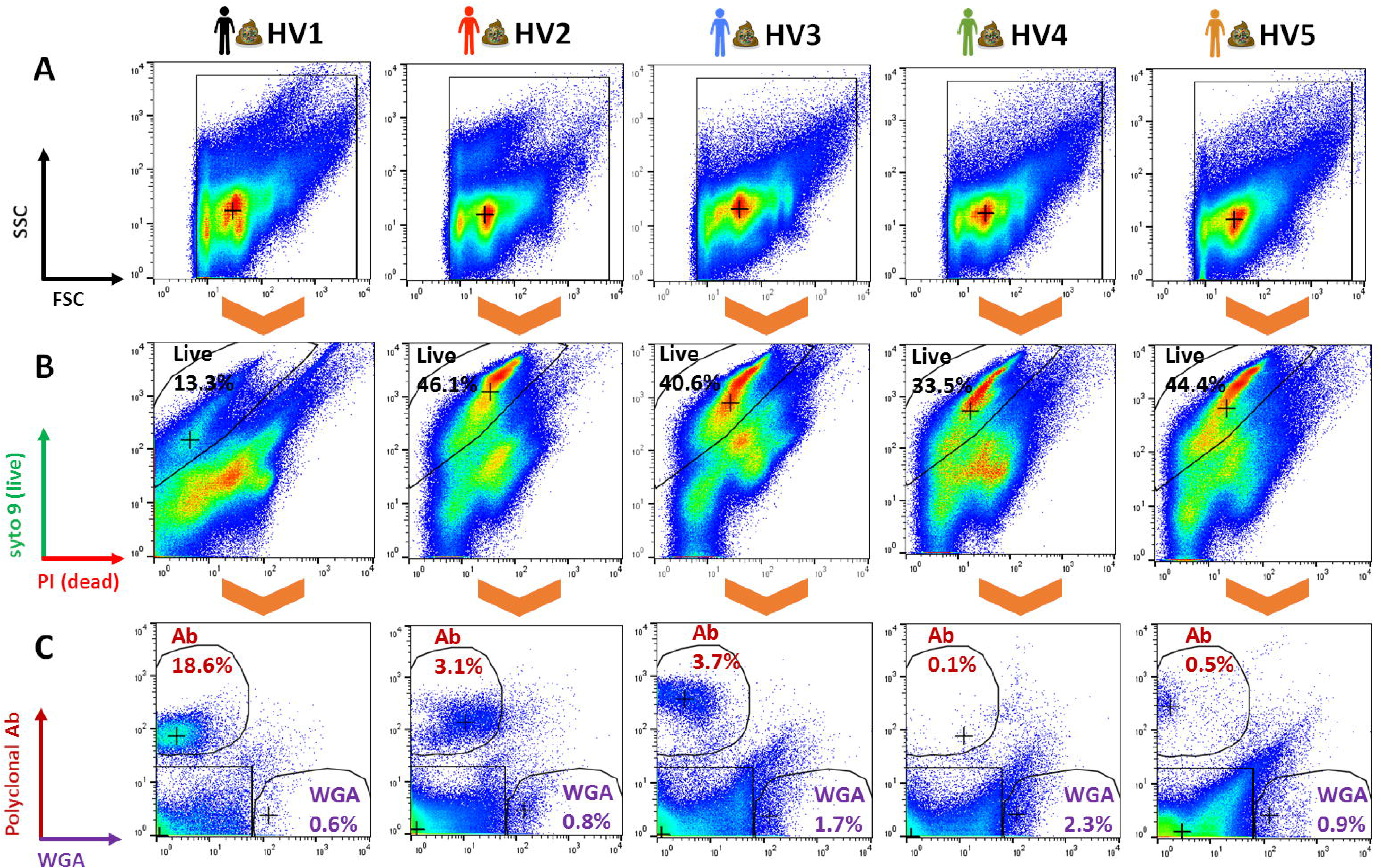
Flow cytometry analysis of fecal microbiota. Fresh fecal samples collected from 5 healthy volunteers were stained using Live/Dead staining (B), polyclonal antibody directed against *F. prausnitzii* ATCC-27766 and ATCC-27768 (C, y-axis), and WGA staining (C, x-axis).

**Table 1.** Proportions of bacteria detected and cultivated using the different gating strategies (Live/Dead, polyclonal antibody and WGA staining), and then identified as putative *F. prausnitzii* strains using species-specific PCR primers. Only antibody-positive or WGA-positive events were sorted for culture.

| Sample | Percentage of live bacteria | Antibodies |  |  | WGA |  |  |
| --- | --- | --- | --- | --- | --- | --- | --- |
|  |  | Percentage | Number of cultivated colonies | Fp PCR-positive | Percentage | Number of cultivated colonies | Fp PCR-positive |
| HV1 | 13.3% | 18.6% | 25 (6.5%) | 0 (0%) | 0.6% | 103 (26.8%) | 6 (5.8%) |
| HV2 | 46.1% | 3.1% | 26 (6.8%) | 5 (19.2%) | 0.8% | 13 (3.4%) | 3 (23.1%) |
| HV3 | 40.6% | 3.7% | 13 (3.4%) | 7 (53.8%) | 1.7% | 100 (26.0%) | 9 (9.0%) |
| HV4 | 33.5% | 0.1% | 61 (15.9%) | 30 (49.2%) | 2.3% | 81 (21.1%) | 1 (1.2%) |
| HV5 | 44.4% | 0.5% | 21 (5.5%) | 6 (28.6%) | 0.9% | 55 (14.3%) | 2 (3.6%) |

**Table 2.** Proportions of bacteria detected and cultivated using the different gating strategies (Live/Dead, polyclonal antibodies and WGA staining), and then identified as putative *F. prausnitzii* strains using species-specific PCR primers. Antibody-positive, WGA-positive and double-positive (antibody+WGA) events were sorted for culture.

| Sample | Percentage of live bacteria | Antibodies |  |  |  | WGA |  |  |  | WGA/Ab |  |  |  |
| --- | --- | --- | --- | --- | --- | --- | --- | --- | --- | --- | --- | --- | --- |
|  |  | Percentage among live | Number of cultivated colonies | Number tested | *Fp* PCR-positive | Percentage among live | Number of cultivated colonies | Number tested | *Fp* PCR-positive | Percentage among live | Number of cultivated colonies | Number tested | *Fp* PCR-positive |
| HV6 | 39.1% | 4.1% | 3 (2.5%) | 2 | 1 (50.0%) | 2.8% | 26 (21.7%) | 12 | 1 (8.3%) | 1.9% | 15 (12.5%) | 3 | 0 |
| HV7 | 61.3% | 0.2% | 20 (16.7%) | 6 | 1 (16.7%) | 1.9% | 29 (24.2%) | 7 | 3 (42.9%) | 0.3% | 12 (10.0%) | 3 | 0 |
| HV8 | 65.9% | 0.1% | 2 (1.7%) | 1 | 0 | 3.7% | 16 (13.3%) | 3 | 1 (33.3%) | 0.4% | 3 (2.5%) | 1 | 0 |
| HV9 | 41.2% | 0.5% | 3 (2.5%) | 2 | 0 | 8.3% | 47 (12.2%) | 5 | 2 (20.0%) | 0.6% | 8 (6.7%) | 3 | 0 |
| HV10 | 52.7% | 4.1% | 18 (15.0%) | 7 | 7 (100.0%) | 3.7% | 16 (13.3%) | 7 | 1 (14.3%) | 1.3% | 26 (21.7%) | 3 | 1 (33.3%) |

## Clonality and phylogenetic analysis of newly isolated *F. prausnitzii* strains

RAPD analysis was used to de-replicate isolates collected from the same donors, resulting in a collection of 9 different strains that were positive with *F. prausnitzii*-specific primers Fprau 02 and Fprau 07 and that presented unique specific RAPD profiles with primers D9355 (Fig. 6). Partial 16S-rRNA encoding genes were amplified and sequenced from these strains, resulting in a collection of 8 confirmed *F. prausnitzii* and 1 Gemmiger-related strains. The phylogenetic tree inferred from these sequences encompassed a variety of strains recovered from the fecal samples of healthy volunteers, with strains belonging to the 3 phylogroups (I, IIa and IIb) described for *F. prausnitzii* [1] (Fig. 7).

**Fig. 6.**
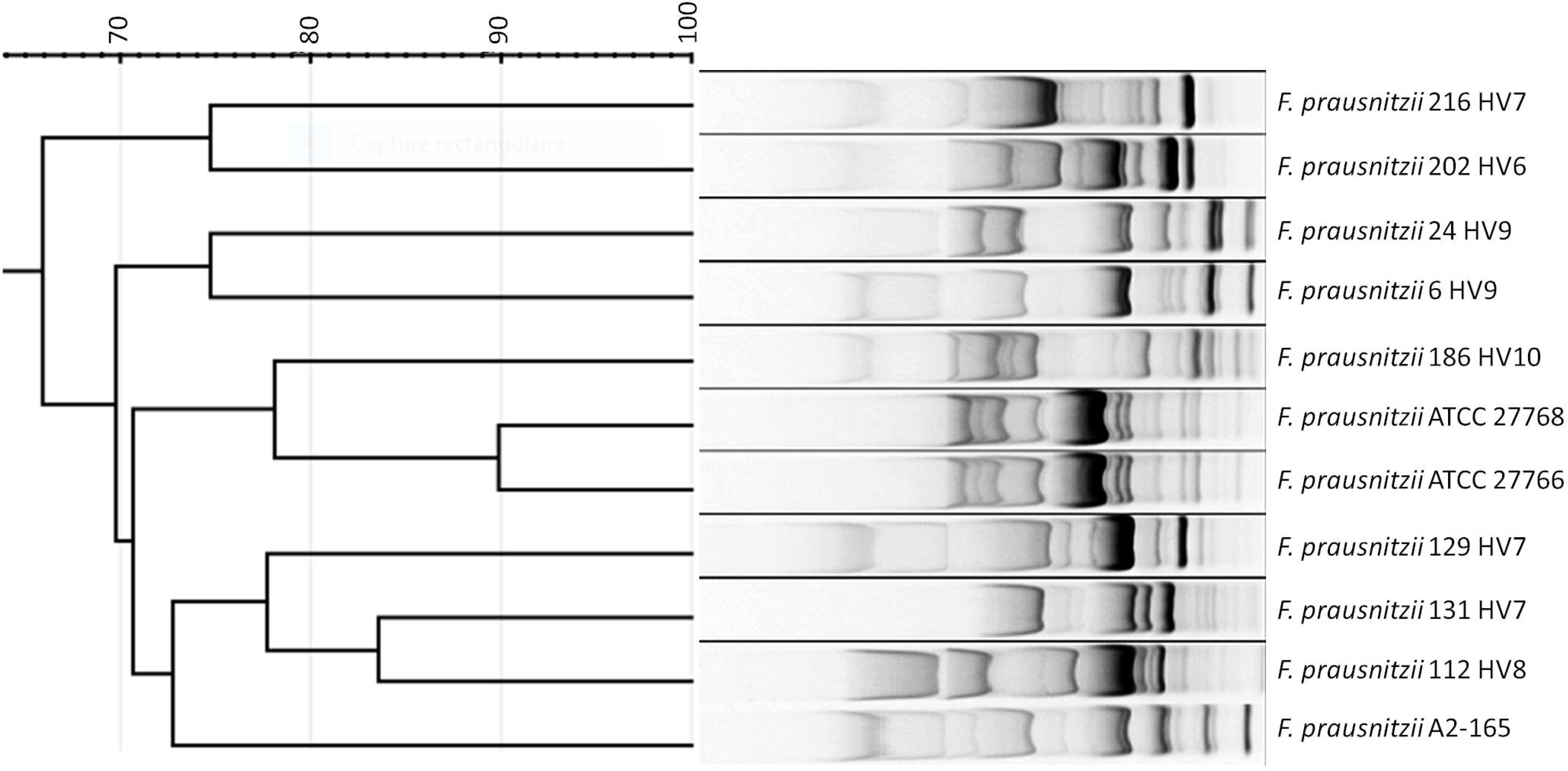
Random Amplified Polymorphism DNA profiles obtained with *F. prausnitzii* reference strains and newly isolated strains using primer D9355.

**Fig. 7.**
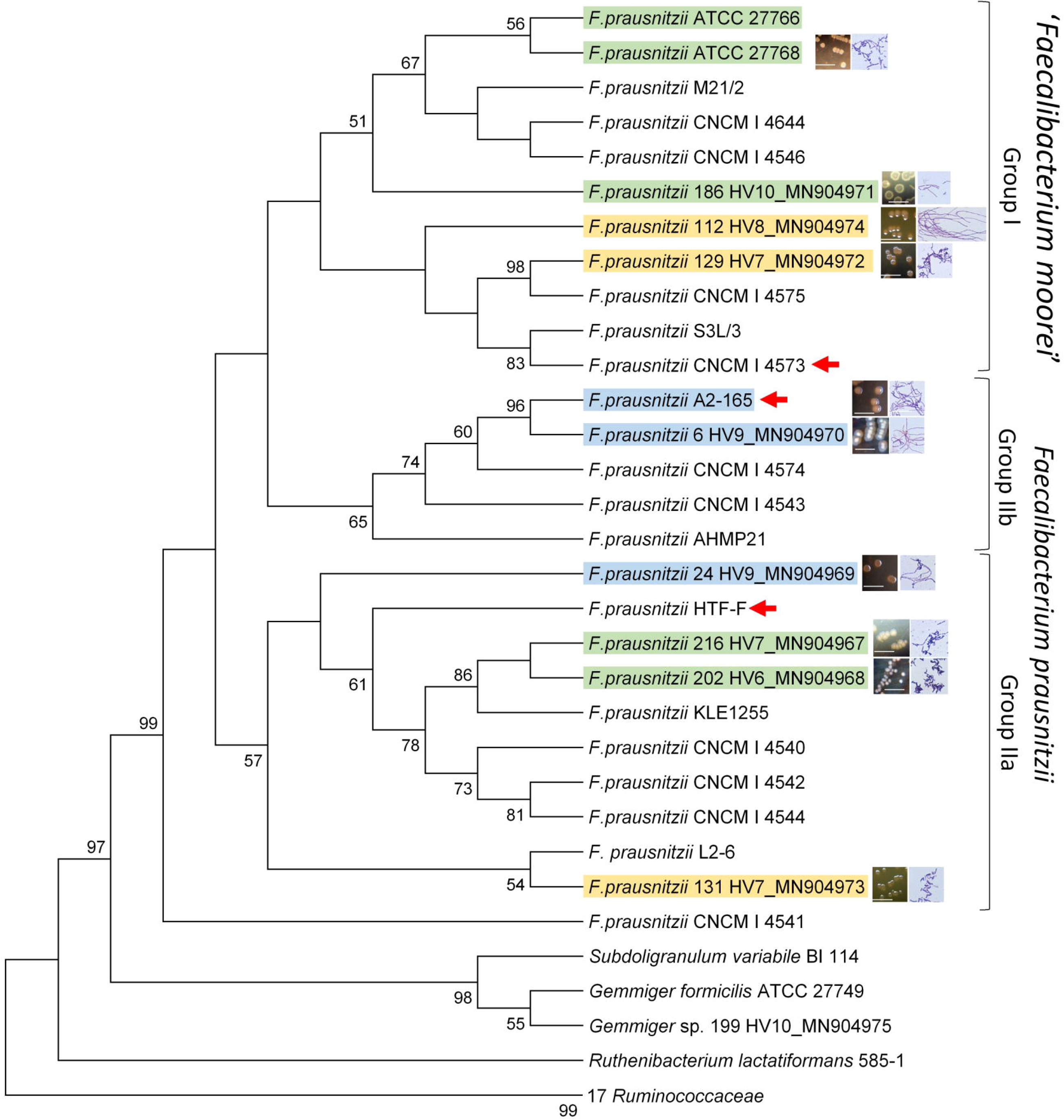
Phylogenetic tree representing newly isolated *F. prausnitzii* strains. The phylogenetic tree was inferred from Muscle alignment of partial 16S rRNA-encoding gene sequences using the Maximum Likelyhood method based on the Kimura 2-parameters model with 1,000 bootstrap replicates. Branch values < 50% are not displayed. The tree was built using reference sequences and outgroups described by [1]. Colonies aspects (bar: 0.5 cm) as well as Gram-stains (100x objective lens) are reported for the strains used in this study. Previously described strains with demonstrated anti-inflammatory activities are indicated with a red arrow. Strains highlighted in blue were efficiently stained with WGA (>90% of the events) but not with the antibodies (<0.5% of the events), strains highlighted in yellow presented moderate WGA staining (approx. 15% of the events) and no reaction with antibodies, and the strains highlighted in green presented variable staining with WGA (10 to 30% of the events) and with the antibodies (30 to 60% of the events).

## Phenotypic analysis of newly isolated *F. prausnitzii* strains

Colonies observed after cultivation on mGAM agar plates showed heterogeneous morphologies. After 72 h incubation some strains appeared as translucent colonies embedded in agar with a fried egg appearance with irregular edges whereas others appeared as tiny bulging colonies. Bacteria appeared as variable length bacilli after Gram staining, with most strains clearly stained as Gram-negative whereas several ones (#202 and 216) appeared as Gram-variable or Gram-positive (Fig. 7). Strain 112 formed long filaments consisting in chains of several bacteria. FCM analysis of the bacteria after double-staining with WGA and with polyclonal antibody confirmed the results observed with the 3 collection strains, with strains n°6 and 24 (highlighted in blue in Fig. 7) being efficiently stained with WGA (>90% of the events) but not with the antibody (<0.5% of the events), strains n°112, 129 and 131 (highlighted in yellow in Fig. 7) presenting moderate WGA staining (approx. 15% of the events) and no reaction with antibody, and the remaining strains (n° 186, 202 and 216, highlighted in green in Fig. 7) presenting variable staining with WGA (10 to 30% of the events) and with the antibody (30 to 60% of the events).

## Evaluation of immunomodulatory potential of the newly isolated *F. prausnitzii* strains

To evaluate the immunomodulatory potential of the newly isolated strains, filtered supernatants of *F. prausnitzii* strains grown for 24 h in Y-BHI medium were added at final concentration of 20% to HT-29 cells cultures in parallel to stimulation with 5 ng/ml TNF-α. Six hours later, the concentration of IL-8 in the cell culture medium was determined using ELISA method. Similarly to previous findings by Martin *et al*. [2], isolates differed in their capacity to reduce IL-8 secretion upon TNF-α stimulation, sometimes showing stronger potential than the reference strains (Fig. 8). Of note, no correlation was found between the final OD_600_ reached by the different strains and their capacity to reduce IL-8 secretion (data not shown).

**Fig. 8.**
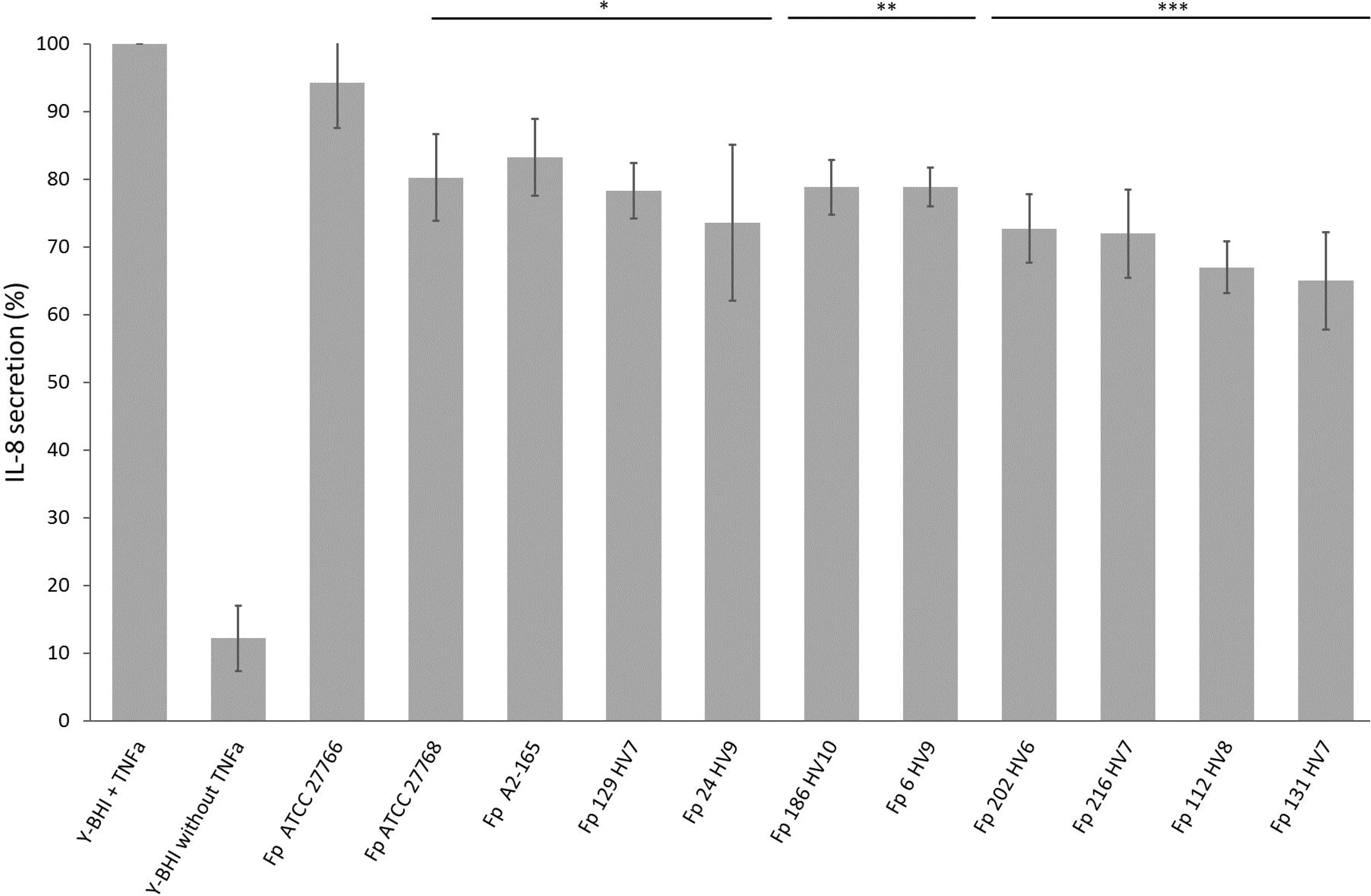
The effect of Faecalibacterium supernatants on IL-8 secretion from HT-29 epithelial cells upon stimulation with 5 ng/mL TNFα. Filtered bacterial culture supernatants were added at final concentration of 20%. Strains that failed to reach the optical density (600 nm) of at least 0.2 were excluded from the experiment. The results are normalized to IL-8 secretion upon stimulation with TNFα and fresh Y-BHI medium and the statistical analysis was made using one-way Anova method comparing each result to this sample. The experiment repeated 5 times so that each strain was tested at least 2 times. * adjusted *P* < 0.05, ** *P* < 0.01, *** *P* < 0.001.

## Discussion

The field of microbiome research has rapidly developed these last few years, with countless associations reported between microbiota composition and specific health conditions. This is especially true for human gut ecosystem, for which dysbiotic microbiota have been associated with metabolic syndrome, inflammatory bowel disease and response to cancer immunotherapy to mention just a few, thus offering new avenues to explore with the ultimate goal to develop new, complementary tools to treat these conditions [3–5]. In this context, collections of bacterial strains are needed in order to explore functionalities and host-bacteria interactions, and go beyond simple association studies. Since many of these commensal species are Extremely Oxygen Sensitive, completely anaerobic conditions are required when applying protocols to cultivate them from fecal material.

FCM analysis of environmental and human-associated microbiota after DAPI-staining have been described, allowing to distinguish specific subgroups and compositional variations in these ecosystems [6–9]. It can be associated with various high-level staining methods including for instance viability staining or Gram-staining, whereas phylogenetically-resolved detection usually involves rRNA-probes that require fixation before hybridization or species-specific antibodies (for a review see [10]). The recent description of FCM and bacterial cell sorting in anaerobic conditions [11] prompted us to explore this technology for ‘targeted culturomics’ of species of interest from fecal material.

As a proof-of-concept study, we chose to focus on the EOS species *F. prausnitzii*. This species represents approximately 5 to 10% of the healthy gut microbiota [12], it has been associated with a number of favorable outcomes in various disease conditions, including lower risk of postoperative recurrence of ileal Crohn’s Disease [13] and improved response to immune check point blockers [14, 15]. Several different mechanisms have been proposed to explain beneficial anti-inflammatory effects, including the production of specific molecules that inhibit the NF-κB pathway [16] or the production of extracellular polymeric matrix that modulates IL-10 and IL-12 production in a TLR-2 dependent way [17, 18]. However anti-inflammatory as well as immunomodulatory effects vary significantly from one strain to another when tested *in vitro* [2]. This is also the case for other important traits such as butyrate production or the production of various enzymes [2]. These differences could partly be explained by the complex phylogeny of the *F. prausnitzii* species that comprise at least 3 different phylogroups, and the probable existence of closely related species [1, 19, 20] that have not been described so far and that were mistakenly identified as ‘*F. prausnitzii*’ in 16S repertoire studies that provide only limited phylogenetic information at the species level. Relative proportions between the different phylogroups in one same individual seem to vary depending on specific disease condition, with phylogroup IIb strains being depleted in Crohn’s Disease patients [21, 22]. It has consequently been proposed to use these relative abundances as a biomarker of disease condition [23].

In our hands, a cell selection process based on specific stains such as Vancomycin BODIPY™-FL was extremely accurate for simple mixtures of bacterial species, which was a prerequisite for experiments with complex microbiota. In addition, the EOS species *F. prausnitzii* sorted after various staining methods remained cultivable and the LIVE/DEAD staining was in good correlation with cultivability, with only very few colonies obtained from PI-stained bacteria. The WGA lectin bound very efficiently to a subset of strains that belong to the phylogroup IIb, whereas strains that belong to the phylogroup IIa and to the phylogroup I (recently proposed as ‘*Faecalibacterium moorei*’ [1]) were less efficiently stained. These differences likely correspond to differences in cell wall compositions, such as for instance the presence of an extracellular polymeric matrix as described for *F. prausnitzii* strain HTF-F [17]. In addition, polyclonal antibody generated by vaccinating rabbits with a mix of the two closely related strains *F. prausnitzii* ATCC 27768 and 27766 that do not bind efficiently WGA were completely unreactive against WGA-stained strains. This could be due to the absence of epitopes, or more likely to extracellular compounds masking epitopes that are exposed in strains that are only weakly or moderately stained by WGA. Whereas these differences correspond to specific phylogenetic groups and/or could possibly be indicative of specific biological properties remains to be explored.

Taking these differences into account, both stains were used to analyze bacteria in fecal samples, and then sort and cultivate antibodies- and WGA-labelled bacteria separately. The LIVE/DEAD staining that was used concomitantly to select bacteria from the live-stained fraction revealed important variations from one sample to another. The sorting strategy resulted in significant proportions of cultivated colonies that were tested positive with *F. prausnitzii*-specific primers, thus confirming the interest of the approach (Tables 1 and 2). Despite high proportion of live-gated bacteria being detected by the antibody in the fecal sample collected from healthy volunteer 1, we were not able to cultivate any *F. prausnitzii* colony from bacteria corresponding to this region of the dot-plot whereas several colonies were obtained from the WGA-positive bacteria. One possible explanation could be that *F. prausnitzii* bacteria detected with the polyclonal antibody require specific nutrients that were not present in our complemented mGAM medium. Co-cultivation with *E. coli* or cultivation in the presence of spent *E. coli* cultivation medium could be tested since it has been reported that *E. coli*-produced quinones are growth factors for *F. prausnitzii* strain KLE1255 [24]. Another explanation could be that this specific strain was not efficiently labelled with the LIVE/DEAD staining, being labelled as ‘live’ whereas it was dead or in a viable but non-culturable state (the proportion of live-stained events was very low for this specific sample).

We choose to use RAPD primer D9355 originally described by Akopyanz *et al* [25] in our strain typing experiments since it generated more complex band profiles compared to short primer 1254 described in the same original paper and that has been previously used for *F. prausnitzii* [26, 27] (data not shown). However primer D9355 did not distinguish between phylogenetically related strains ATCC 27766 and ATCC 27768. Keeping these limitations in mind, RAPD was still a valuable tool when used with DNA purification using Instagen kit for the rapid typing of PCR-positive colonies identified from the same original fecal sample. Interestingly, isolates 6 and 24 that were cultivated from the same original fecal sample presented similar RAPD profiles. However they were classified as two different strains due to several slight differences. Whether these differences reflect evolution of one commensal strain through gain and/or loss or several genes [28] or should be attributed to technical biases will be confirmed when sequencing the complete genomes.

It was shown previously that *F. prausnitzii* secrets factors capable of modulating the response of the host cells to inflammatory stimuli [2, 13]. Intestinal epithelial cells are the first to interact with lumen-resident microbiota and their products, regulating the downstream inflammatory outcome. In this way, a presence of supernatant from *F. prausnitzii* was shown to decrease IL-8 secretion by epithelial cells (HT-29) in response to TNFα stimulation [29]. Similar results were observed in our experiments, with marked differences between strains confirming the need to take into account these phenotypic variations when choosing one specific strain for NGP development. These characterizations should be completed with the exploration of additional phenotypic traits that can be important for host-bacteria interactions such as the induction of IL-10 secretion, resistance to acidic pH, adhesion to mucine, Short Chain Fatty Acids production *etc* [2].

As a conclusion, this proof-of-concept study confirms that FCM is well adapted for complex bacterial microbiota studies. When used in conjunction with appropriate staining, it gives a general overview of microbiota composition and variations in longitudinal studies [6], including bacterial load which is an important piece of information [30]. In addition, the use of more specific staining such as antibodies is a promising strategy to target, sort and cultivate species of interest from these complex ecosystems. Recent studies demonstrate that these antibodies can be generated using a reverse genomics approach [31], which opens important avenues since approximately 70% of the gut microbial species still lack cultured representatives [32].

This should be accompanied by specific technological developments in the field of FCM to allow simple, commercially-available solutions enabling routine sorting experiments in controlled atmosphere conditions, which will be of strong interest for commensal bacteria but also for cellular biology applications necessitating oxygen conditions that are close to in vivo conditions [33, 34].

## Methods

### Bacterial strains and growth conditions

The following reference strains were used in this study: *Faecalibacterium prausnitzii* strains DSM-17677 (A2-165), ATCC-27766 and ATCC-27768, *Escherichia coli* ATCC 35218 and *Enterococcus faecalis* ATCC 29212. *F. prausnitzii* strains were cultivated using modified Gifu Anaerobic Medium (mGAM, HyServe 05426) complemented with 30% bovine rumen, cellobiose (1 mg/ml) and inulin (1 mg/ml) with (plates) or without (broth) 1.5% agar. All media and reagents were reduced for at least 48 h in the BACTRON600 (Sheldon) anaerobic chamber before use, and cultures were incubated at 37 °C in the chamber. *E. coli* was cultivated in LB medium and *E. faecalis* in BHI medium, both under aerobic condition at 37 °C.

### Polyclonal antibodies

Rabbit polyclonal antibodies (pAb) were produced in New Zealand rabbits using a standard 53 days protocol (Covalab). Rabbits were immunized with a 50/50 mix of phylogenetically related strains *F. prausnitzii* ATCC-27766 and ATCC-27768. Briefly, rabbits were first immunized with two injections of 1 × 10^8^ heat-killed (80 °C / 30 min) bacteria mixed with complete Freund’s adjuvant (1:1 v:v), then boosted with a third injection of 2 × 10^8^ of the same heat-killed bacteria. After rabbit bleeding, sera were harvested and IgGs were purified on protein A and labeled with Alexa Fluor™ 647 using the protein labeling kit (Thermo Fisher Scientific) as recommended by the manufacturer.

### Non-specific staining procedures

Wheat germ agglutinin (WGA-Alexa Fluor™ 647 Conjugate or WGA-CF405M Conjugate, Biotium) staining was performed as previously described [35] with slight modifications. Briefly, bacteria were washed once in phosphate buffer saline (PBS), then numbered by flow cytometry (FCM) using counting beads and adjusted at 1 × 10^6^ events/ml in staining solution (PBS, WGA-647 30 μg/ml). Bacteria were incubated for 20 min at room temperature in the dark. Vancomycin BODIPY™-FL (Thermo Fisher Scientific) staining was performed as for WGA staining, with vancomycin BODIPY™-FL being used at the final concentration of 2 μg/ml. Live/Dead staining was performed using the LIVE/DEAD BacLight Bacterial Viability kit (SYTO 9/PI, Thermo Fisher Scientific) as recommended by the manufacturer. After WGA, Vancomycin BODIPY™-FL or LIVE/DEAD staining, bacteria were washed in PBS and then analyzed within 30 min using an Influx^®^ (Becton-Dickinson) cell sorter equipped with a 200 mW-488 nm laser, a 120 mW-640 nm laser and a 100 mW-405 nm laser.

### Validation of Gram staining and sorting precision

Bacterial cell sorting was first explored with facultative anaerobes to check sorting precision and live bacteria recovery. Gram-negative (*E. coli* ATCC 35218) and Gram-positive (*E. faecalis* ATCC 29212) bacteria were cultivated overnight in BHI broth and then numbered using counting beads. They were adjusted to a concentration of 1 × 10^6^ bacteria / ml and equally mixed before labeling with the SYTO 62 DNA staining dye (Thermo Fisher Scientific) at 1 μM and the Gram-positive dye Vancomycin BODIPY™-FL at 2 μg/ml for 15 mn. Bacteria were interrogated with a 488-nm laser and a 640-nm laser and SSC and FSC parameters were collected for each cell, with FSC as the data collection trigger. Sorting gates were defined based on 540/30 nm and 670/30 nm fluorescences. Single cells were sorted into 16 × 24 matrices onto URI*Select*™ 4 agar plates (Bio-Rad) prepared in Nunc™ OmniTray™ Single-Well Plates (Thermo Scientific). The first two columns (Fig.1 B-1) were inoculated with single cells corresponding to Vancomycin BODIPY™-FL unlabeled gate. The next two columns (Fig.1 B–2) were inoculated with single cells corresponding to the Vancomycin BODIPY™-FL labeled gate. Then two columns were first inoculated with Vancomycin BODIPY™-FL-negative single cells and then with Vancomycin BODIPY™-FL-positive single cells (Fig.1 B–3). Remaining columns (Fig.1 B–4) were inoculated with single cells randomly collected from the gate designed from SSC/FSC representation and encompassing both *E. coli* and *E. faecalis* bacteria.

### Anaerobic sorting of single strains

The BD Influx^®^ cell sorter used for anaerobic sorting has been described before [11]. Briefly, a glove box was plugged on the sorting chamber and was then fed with nitrogen to deplete oxygen from the sorting chamber. Oxygen concentration in the glove box was monitored using a Fibox 4 trace detector (PreSens). For sorting experiments, reduced mGAM plates were transferred from the anaerobic chamber to the cell sorter glove box using sealed bags. Nitrogen was then injected and anaerobic sorting experiments were started when the oxygen concentration was measured below 0.7%. Analysis and sorting followed by cultivation of stained and unstained bacteria was used to evaluate the impact of the process on *F. prausnitzii* cultivability. Bacteria used for sorting experiments were anaerobically cultivated for 48 h at 37°C on mGAM plates. One colony was then sub-cultivated in mGAM broth for 24 h at 37°C. Bacteria were then washed in reduced PBS before staining with LIVE/DEAD, WGA-647 or Vancomycin BODIPY™-FL as described above. After washing, stained bacteria were suspended in reduced PBS containing 0.5 mg/l resazurin, 2.1 mM soldium sulfure and 2.8 mM L-cystein HCl, and the suspensions were covered with 500 μl of paraffin oil to prevent oxygen exposure. The tubes were taken out of the anaerobic chamber and bacteria were analyzed and sorted within 30 min. Sorting speed was adjusted at 1,000 events per second and four series of 1, 3, 10, 30, 100, 300 or 1,000 events were sorted on one single spot for each tested condition. Once sorting experiments were achieved, plates were re-introduced in sealed bags and transferred in the anaerobic chamber where they were incubated at 37 °C for 2 to 3 days before observation. Experiments were repeated at least 3 times for each strain.

### Anaerobic sorting and cultivation from fecal material

Fecal samples were collected at home by healthy volunteers and immediately transferred in plastic pouches that were then sealed after adding an AnaeroGen™ sachet (Oxoid). Samples were shipped to the lab within 2 h at ambient temperature. All subsequent steps were performed in the anaerobic chamber. For each sample, 1 g of fecal material was suspended in 10 ml PBS and homogenized by vortexing using 2.4 mm glass beads. Suspensions were then 10-fold diluted in PBS and filtered through a 70-μm cell strainer (Biologix). After an additional 100-fold dilution in PBS, bacteria were numbered by FCM using counting beads (LIVE/DEAD™ BacLight™ Bacterial Viability and Counting Kit, Thermo Fisher Scientific). LIVE/DEAD staining was used to select live bacteria, and polyclonal antibodies conjugated with Alexa Fluor™ 647 were used in combination with WGA-CF405M to gate *F. prausnitzii* bacteria. Labelling was performed in anaerobic conditions for 20 min in the dark, and then bacteria were washed in reduced PBS before analysis. After washing, stained bacteria were suspended in reduced PBS and the suspensions were covered with 500 μl of paraffin oil to prevent oxygen exposure. The tubes were taken out of the anaerobic chamber and bacteria were analyzed and sorted within 30 min. Bacteria were gated based on FSC/SSC parameters. Among them, live ones were selected according to SYTO 9/PI fluorescence. Two additional gates were then defined for the first series of sorting experiments: gate A encompassing antibody-positive/WGA-negative events and gate B for the antibody-negative/WGA-positive events. 384 events collected from each of these two gates were sorted on mGAM plates. In the second series of experiments, a third gate (gate C) was defined for the double positive population, and 120 events were sorted in anaerobic conditions onto mGAM plates for each of the 3 gated regions. Plates were then incubated for 5 days at 37°C in anaerobic conditions. Several colonies with morphologies compatible with typical *F. prausnitzii* colonies (*i.e*. small, flat colonies encrusted in the agar plates) were then selected for screening by PCR using previously described specific primers Fprau 02 (5’-GAG CCT CAG CGT CAG TTG GT-3’) and Fprau 07 (5’-CCA TGA ATT GCC TTC AAA ACT GTT-3’) [13], and further identified by sequencing the almost complete 16S rRNA-encoding genes (GenBank accession numbers: MN904967 to MN904975).

### Clonality and phylogenetic analysis of newly isolated *F. prausnitzii* strains

Random Amplified Polymorphic DNA (RAPD) analysis was conducted to distinguish strains. Briefly, DNA was isolated from 100 μL of fresh overnight Y-BHI cultures of *F. prausnitzii* strains using InstaGene™ Matrix (BioRad) kit according to the manufacturer guidelines. This DNA preparation served as a template for PCR reaction using primer D9355 and a PCR program described by Akopyanz et al. [25]. To estimate the phylogenetic distances, images of the agarose gels taken upon electrophoresis of the PCR products were analyzed using GelJ_v2 software [36] using Jaccard similarity method and UPGMA linkage with a tolerance value of 1. For phylogenetic analysis, 16S rRNA-encoding genes were aligned using Muscle [37] integrated in MEGA7 [38] with default parameters. The phylogenetic tree was inferred using the Maximum Likelyhood method based on the Kimura 2-parameters model with 1,000 bootstrap replicates [2].

### Primary phenotypic characterization of newly isolated *F. prausnitzii* strains

Bacteria were cultivated on mGAM plates for 24 to 48 h at 37 °C in anaerobic conditions for the documentation of colony morphology. Gram-staining was performed with bacteria cultivated for 48 hours in mGAM broth. Staining with polyclonal antibody conjugated to Alexa Fluor™ 647 and with WGA-CF405M conjugate were conducted as described above. Interleukin-8 (IL-8) secretion assay was done according to the procedure described by Kechaou *et al*. [39]. Briefly, 7 days before the experiment 50,000 HT-29 cells per well were seeded in 24-well culture plates in 0.5 ml of DMEM+Glutamax medium (Gibco) supplemented with 10% fetal bovine serum (FBS). 24 h before the experiment, the culture medium was changed for a 0.4 ml medium with 5% heat-inactivated FBS. On the day of experiment, fresh cultures of *F. prausnitzii* grown for 24 h were centrifugated for 5 min at 5,000 x g and the supernatant was filtered through 0.22 μm filters (Merck). Then 100 μl of the cleared bacterial supernatants (or fresh Y-BHI medium alone) were added to the cell cultures (final volume of 20%). Cells were stimulated simultaneously with human TNF-α (5 ng/ml, Peprotech) for 6 h at 37°C in 10% CO_2_. All samples were analyzed in triplicates. After co-incubation, cell supernatants were collected and stocked at −80°C until further analysis of IL-8 concentrations by ELISA (R&D Systems) according to manufacturer’s guidelines. The results were normalized using total protein content of the cells that was determined using Pierce™ BCA Protein Assay Kit (Thermo Fisher Scientific). Experiments have been performed in triplicates in five independent experiments.

## Declarations

### Ethics approval and consent to participate

Stools samples were anonymously collected from healthy volunteers. As required by the French regulation, this collection was declared to the French ministry of Higher Education, Research and Innovation (n°DC-2015-2513) and received an approval by the ethical committee in March 2016. Each volunteer received an information and signed a consent before participating to the study.

### Consent for publication

Not applicable.

### Availability of data and material

16S-rRNA encoding gene sequences of the new isolates described in this study are available under GenBank accession numbers MN904967 to MN904975.

### Authors’ contributions

VT and SB conceptualized and designed the experiments. JB contributed to the technical setup of the anoxic cell sorter. MN, AD, MA and SB performed the staining and sorting experiments, and contributed to strains identification using specific PCR and 16S rRNA gene sequencing. IB performed the RAPD and IL-8 secretion assay. SB and VT wrote the manuscript. All authors read and approved the final manuscript.

## Acknowledgements

Not applicable.

